# Habenula-ventral tegmental area functional coupling and risk-aversion in humans

**DOI:** 10.1101/2024.11.01.621507

**Authors:** Wanjun Lin, Jiahua Xu, Xiaoying Zhang, Raymond J Dolan

**Affiliations:** Max Planck University College London Centre for Computational Psychiatry and Ageing Research, University College London, Queen Square Institute of Neurology, London, UK; Psychiatry Research Center, Beijing Huilongguan Hospital, Peking University Huilonguan Clinical Medical School, Beijing, China; State Key Laboratory of Cognitive Neuroscience and Learning, IDG/McGovern Institute for Brain Research, Beijing Normal University, Beijing, China; Wellcome Centre for Human Neuroimaging, University College London, Queen Square Institute of Neurology, London, UK

## Abstract

Maladaptive responses to uncertainty, including excessive risk avoidance, are linked to a range of mental disorders. One expression of these is a pro-variance bias (PVB), wherein risk-seeking manifests in a preference for choosing options with higher variances/uncertainty. Here, using a magnitude learning task, we provide a behavioural and neural account of PVB in humans. We show that individual differences in PVB are captured by a computational model that includes asymmetric learning rates, allowing differential learning from positive prediction errors (PPEs) and negative prediction errors (NPEs). Using high-resolution 7T functional magnetic resonance imaging (fMRI), we identify distinct neural responses to PPEs and NPEs in value-sensitive regions including habenula (Hb), ventral tegmental area (VTA), nucleus accumbens (NAcc), and ventral medial prefrontal cortex (vmPFC). Prediction error signals in NAcc and vmPFC were boosted for high variance options. NPEs responses in NAcc were associated with a negative bias in learning rates linked to a stronger negative Hb-VTA functional coupling during NPE encoding. A mediation analysis revealed this coupling influenced NAcc responses to NPEs via an impact on learning rates. These findings implicate Hb-VTA coupling in the emergence of risk preferences during learning, with implications for psychopathology.

## Introduction

Decision-making under uncertainty is a daily challenge, with maladaptive responses proposed as a mechanistic account for a range of psychopathologies, including anxiety (1, 2). depression (3, 4) and gambling disorders (5). More specifically, risk aversion manifesting as a disposition to prefer an option with lower uncertainty/variance (6), is linked to higher anxiety and depression scores in the general population (7). This renders it important to understand how risk-related biases emerge as individuals deal with an uncertain environment.

We developed a simple magnitude learning task (8), wherein participants learn and choose between options with different value distributions, as detailed in Fig. 1 (varying means and/or variances). This task enables manipulation of distribution variance/uncertainty independently from its mean, in contrast to probabilistic learning tasks where risk uncertainty and value are related (9). Previous studies show that when presented with two options, drawn from value distributions with the same mean but different variances, human and non-human primates show a preference for an option with the higher variance, the very opposite to risk aversion, and known as “pro-variance bias” (PVB) (10, 11).

**Fig. 1.**
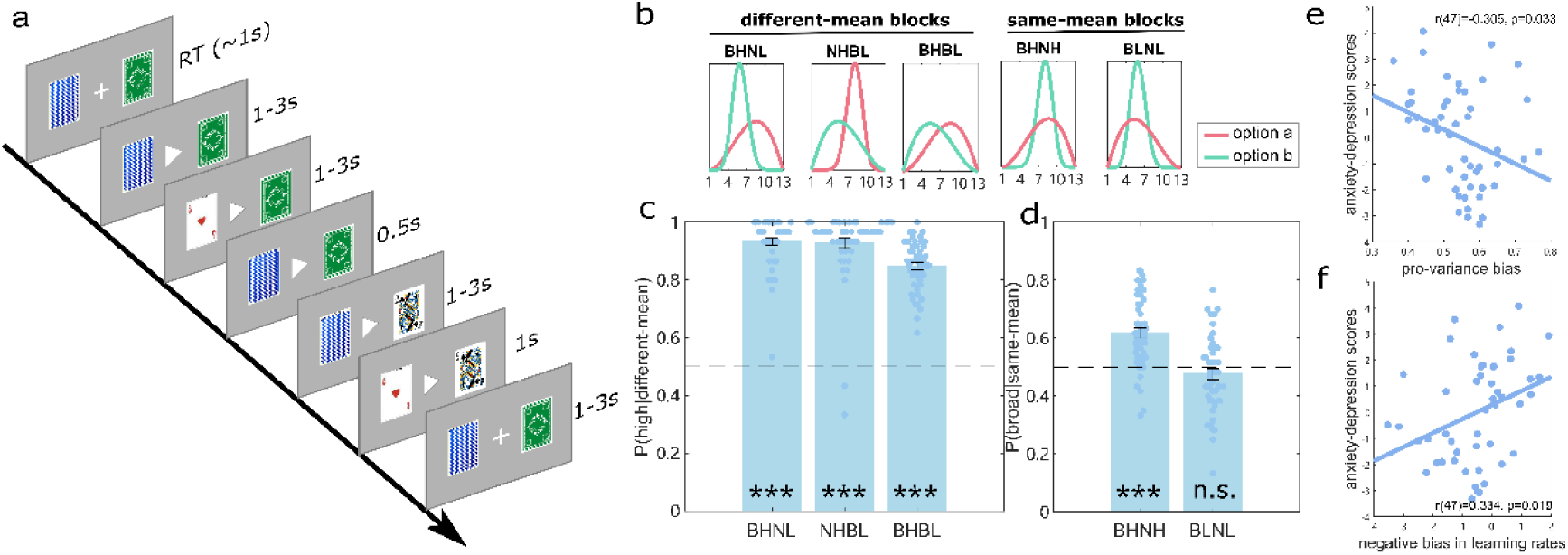
Experimental design. **a)** task structure. Within a task block, participants were presented with the same two card decks across 30 trials and asked to choose from one of the decks. After entering a choice, a triangle in the centre of the screen pointed to the chosen deck. Following this, a card from each deck was shown sequentially to participants (1-3s), with the order of presentation counter-balanced across trials, with the two cards then shown together for 0.5s. **b)** Block types. The two card decks in a block could have the same or different means (high or low), and the same or different variances (narrow or broad). This entailed eight blocks in all comprising four different-mean blocks (one broad-high vs narrow-low (BHNL) block, one narrow-high vs broad-low (NHBL), and two broad-high vs broad-low (BHBL) blocks) and four same-mean blocks (two broad-high vs narrow-high (BHNH) and two broad-low vs narrow-low (BLNL)). **c)** The percentage of choosing the higher option for the different-mean blocks, with dots (light blue) representing data points for each participant for that block type. **d)** The percentage of choosing the broader option for the same-mean blocks. The dots represent data points for each participant for that block type. **e)** PVB negatively correlated with anxiety-depression scores. **f)** model-derived negative bias in learning rates correlated with the anxiety-depression scores. Error bars indicate standard errors (s.e.); *p < 0.05; **p < 0.01; ***p < 0.001.

Previously we showed, both in simulations and empirical data, that a modified Rescorla-Wagner (RW) learning model (12), i.e. a two-learning-rates RW model (2lr-RW), captured individual differences in a PVB (8). In brief, this 2lr-RW model incorporated separate learning rates for positive (PPEs) and negative (NPEs) prediction errors (a better or worse outcome relative to expected values (EVs) respectively), allowing EVs to differ from the true outcome means. Thus, a positive-biased agent, with a higher positive compared to negative learning rate, will integrate more PPEs than NPEs into their EVs compared to NPEs during learning leading to enhanced EVs in a higher variance environment and the emergence of a PVB, i.e. risk-seeking. We note evidence from congenital learned helpless (cLH) rats, who manifest a negative learning bias, also show less risky choices in an entirely different task (13). In the current study, we ask which brain regions support such bias in learning rates that give rise to individual differences in risk preferences.

Dopamine (DA) in particular is linked to risk behaviour as evidenced by an impact of dopaminergic medication on risk-related decision-making (14–16). Optogenetic activation of the midbrain DA neurons has provided evidence for a causal influence on risky decisions (17, 18). DA neurons in the ventral tegmental area (VTA) have a central role in motivation and learning (19, 20) by signaling reward prediction errors (RPEs) via increased activity for unexpected rewards and decreased activity for unexpected reward omissions (21, 22). More recently, the lateral habenula (Hb) has been shown to negatively mirror DA RPEs (23–25). Moreover, stimulation of Hb neurons inhibits midbrain DA neurons (23), suggesting lateral Hb is a source of NPE signaling to DA neurons. Critically, lesioning the lateral Hb attenuates an inhibition of the VTA DA neuron activity seen in response to reward omissions (26), resulting in a decrease in negative learning rates relative to positive learning rates (i.e. a positive learning bias) (26). On this basis, we specifically ask whether BOLD responses to PPEs and NPEs in human VTA and Hb, and their functional connectivity, relate to a learning rate bias estimated using our magnitude learning task.

VTA DAergic neurons have strong projections to nucleus accumbens (NAcc) and medial prefrontal cortex (mPFC) (27, 28), with both regions expressing robust RPEs signals (29, 30). At a behavioural level, phasic stimulation of NAcc dopamine receptor type-2(D2R)-expressing cells increases risk-aversion, while pharmacological inactivation of mPFC increases risky choices (31, 32). On this basis, we were interested in examining whether activity in downstream dopaminergic regions (NAcc and mPFC) support individual differences in PVB. Additionally, a recent study indicates that NAcc encodes variance in outcomes, consistent with a suggestion that NAcc supports a representation of reward distribution (33). Similarly, it has been shown that mPFC computes state uncertainty (34) as well as encoding a reward distribution (35).

In the current study, we investigate the neural mechanisms underlying individual differences in PVB. Using a bespoke magnitude learning task (Lin and Dolan 2024), combined with partial field-of-view (FOV) ultra-high field (7T) fMRI, we examined whether a priori regions of interest (ROIs) that included Hb, VTA, NAcc and mPFC encode trial-by-trial PPEs and NPEs. Critically, we also asked whether functional coupling between regions that support task-related value-updating relates to individual differences in PVB.

## Results

### Lower Pro-variance bias is associated with higher anxiety and depression

57 participants recruited from general population performed a magnitude learning task (see Fig. 1) whilst inside a 7T scanner (8 were excluded from analyses based on pre-specified criteria (see Methods for details)). To introduce variation in PVB we recruited participants with a spectrum of anxiety and depression scores, features previously linked to individual differences in risky decision-making (7). Participants chose between two card decks associated with different value distributions, wherein we independently manipulated the variance and mean of the distributions (Fig. 1 b). Critically, in half the blocks, participants were presented with options with identical means but different variances (the same-mean blocks, see Fig. 1 b), enabling indexing of how choices were impacted by variance.

We found that, for different-mean blocks, the choice percentage for the higher mean option was significantly above chance level (0.5) (see Fig. 1 b&c, all t(48)>24.111, p<.001). As expected, participants performed less well in the BHBL (both options with high variance) blocks compared to blocks where one of the distributions was narrow (i.e. the NHBL and BHNL block) (both t(48)<-4.020, p<.001). These data indicate participants learned the task, with higher uncertainty impacting learning a value difference.

We next examined whether, for blocks with the same mean, there was a preference for the broader options. Consistent with previous findings (8), participants on average preferred the broader over the narrower option in the both-high (BHNH) blocks (t(48)=6.717, p<.001). However, for the both-low (BLNL) blocks the percentage of choosing the broader option did not differ from chance level (t(48)=-1.215, p=.230). These results suggest participants manifest a pro-variance bias, particularly for the both-high blocks.

Next, we asked if individual differences in pro-variance bias (mean over all the same-mean blocks) were associated with task-independent measures of individual differences in self-reported anxiety and depression. We found a pro-variance bias was negatively correlated with State-Trait Anxiety Inventory questionnaire scores (STAI) (r(47)=-.314, p=.028) and marginally with the Zung self-rating depression scale (SDS) (r(47)=-.280,p=.051). These results replicated findings from an independent online pilot study (n=213): r(211)=.174, p=.011 for the STAI, and r(211)=-.185, p=.007 for the SDS (see Fig. S1 & supplementary method). To simplify the presentation, we provide overall scores for anxiety and depression (Anxiety-Depression scores) by summing the z scores of the SDS and STAI, see Fig. 1 e, r(47)=-0.305, p=.033).

Pro-variance bias could arise from a lack of symmetry in integrating positive and negative prediction errors into prior expectations. Using computational modelling we found that, compared to a suite of alternatives, a 2lr-RW was the best-fitting model (Fig. S2a, see details in Methods & supplementary). Consistent with a hypothesis of a learning bias, model-derived negative biases in learning rates (the ratio of the negative learning rate to the sum of both learning rates, see Method) accounted for individual differences in PVB (r(47)= -.840,p<.001), and correlated with Anxiety-Depression scores (r(47)=-.334, p=.019, Fig. 1 f). This suggests that negatively biased learning is a potential computational mechanism underlying an attenuation of a pro-variance bias (or increased risk avoidance), particularly in people with higher anxiety and depression trait scores.

### Habenula encodes negative prediction errors in humans

We next examined a candidate neural basis for the observed PVB by first ascertaining the expression of PPEs and NPEs within a priori regions of interest, namely the Hb, VTA, NAcc and mPFC. Here we used anatomical masks [28] for NAcc and VTA, and a functionally defined ROI for the ventral medial prefrontal cortex (vmPFC) as derived from a meta-analysis of value-based decision-making tasks (see Fig. 2 b-d) (36). Due to the relatively small size of the Hb (around 30 mm^3^ in volume in humans) (37), and to derive a more precise study-based Hb ROI based on the subjects’ ultrahigh-resolution T1 images (0.7mm^3^), we manually drew an anatomical Hb mask in each participant for each hemisphere (see Methods for more details). We ran a model-based fMRI analysis with trial-by-trial PPEs and NPEs for the chosen option calculated using the best-fitted parameters for each participant from the 2lr-RW model (see Computational modelling and GLM 1 in Methods).

**Fig. 2.**
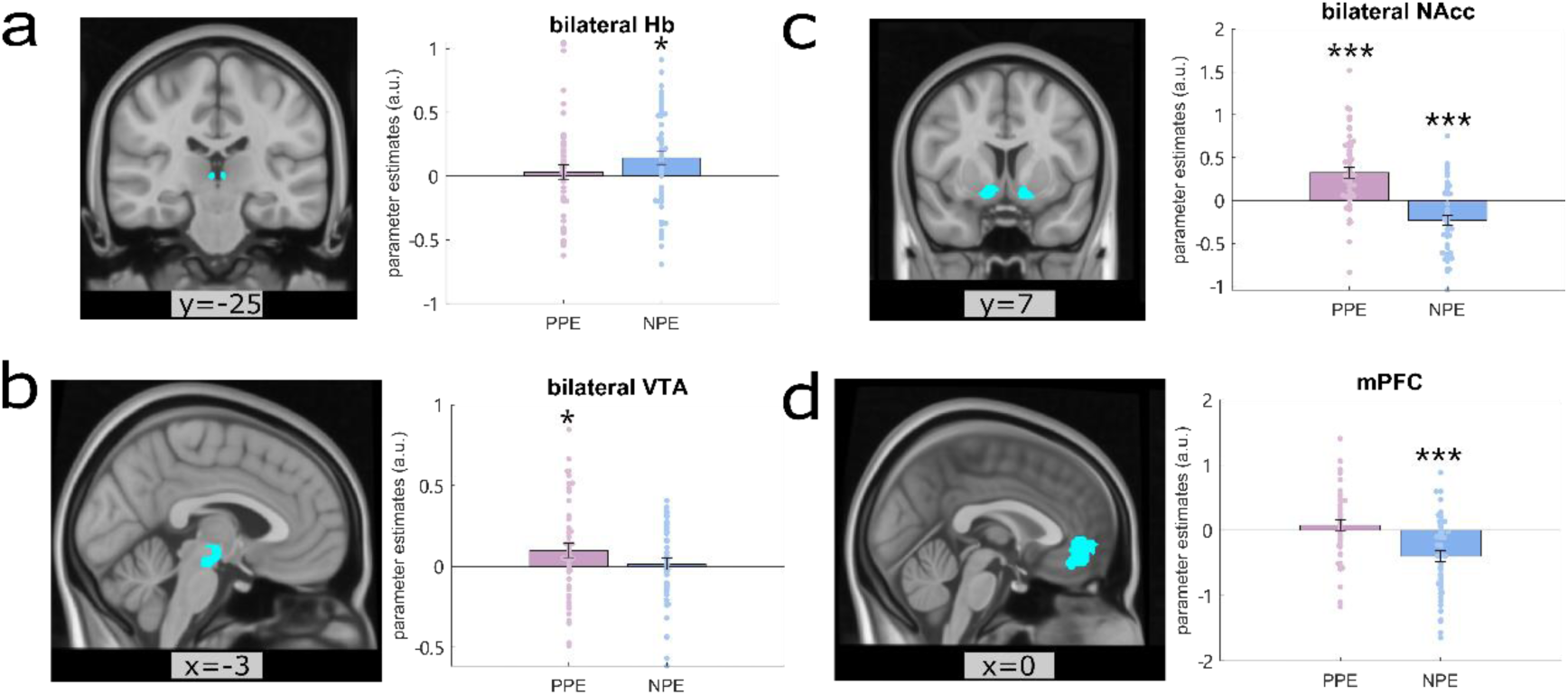
Regions encode prediction errors. **a)** BOLD responses in the bilateral habenula (Hb) were positively modulated by negative prediction errors (NPEs) (p=.016), but not by positive prediction errors (PPEs) (p=.571). Left: a coronal view of the bilateral Hb anatomical mask. Right: parameter estimates of BOLD responses to PPE and NPE, respectively, within bilateral Hb anatomical mask. **b)** BOLD responses in the bilateral ventral tegmental area (VTA) showing positive modulation to positive prediction errors (PPEs) (p=.028) but not to negative prediction errors (NPE) (p=.672). Left: a sagittal view of VTA anatomical mask. Right: parameter estimates of BOLD responses to PPE and NPE, respectively, in this bilateral VTA anatomical mask. **c)** BOLD responses in bilateral nucleus accumbens (NAcc) were positively modulated by PPE and negatively by NPE (both p<.001). Left: a coronal view of the bilateral NAcc anatomical mask. Right: parameter estimates of BOLD responses to PPE and NPE in this bilateral NAcc anatomical mask. **d)** BOLD responses in the ventral medial prefrontal cortex (vmPFC) were negatively modulated by NPE (p<.001) but not PPE (p=.348). Left: a sagittal view of the vmPFC functional defined mask. Right parameter estimates of BOLD responses to PPE and NPE, respectively, in this vmPFC mask. Each dot on the bar graph represents a parameter estimate for each participant. Error bars indicate standard errors (s.e.); *p < 0.05; **p < 0.01; ***p < 0.001.

At choice outcome, VTA BOLD responses were positively modulated by trial-by-trial PPEs (t(48)=2.261, p=.028, see Fig. 2b), but not by NPEs (p=.672). By contrast, NAcc BOLD responses were modulated by both positive and negative prediction errors (Fig. 2c), involving enhanced activation to PPEs (t(48)=5.028, p<.001) and a relative de-activation to NPEs(t(48)=-3.886, p<.001). The vmPFC showed a relative de-activation as a function of NPEs (t(48)=-4.651, p<.001, Fig. 2d) but no response to PPEs (p=.348). Whole-FOV (partial FOV data) group-level results for the NPEs and PPEs accorded with these ROI results (see Fig. S4).

In bilateral Hb, combining left and right Hb masks, we found a positive response to NPEs (see Fig. 2a, t(48)=2.494, p=.016) but not to PPEs (p=.571). A broadly similar pattern of results was evident when we examined the left (NPE: t(48)=1.965, p=.055) and right Hb (NPE: t(48)=2.223, p=.031) alone (see Fig.S5). The overall pattern of results is consistent with habenula encoding negative prediction errors (NPEs), where worse-than-expected outcomes engender a stronger BOLD signal.

We next asked whether BOLD responses to PPE and NPE in target ROIs (i.e. VTA, NAcc, Hb and vmPFC) related to a bias in learning rates. We found a marginally significant negative association between NAcc responses to NPEs and a negative bias in learning rates (see **Fig. 3** b, r(47)=-.274, p=.057). While this suggests NAcc encoding of negative prediction errors relates to a greater negative learning bias, we caution the effect is not strong and below a conventional significance level.

**Fig. 3.**
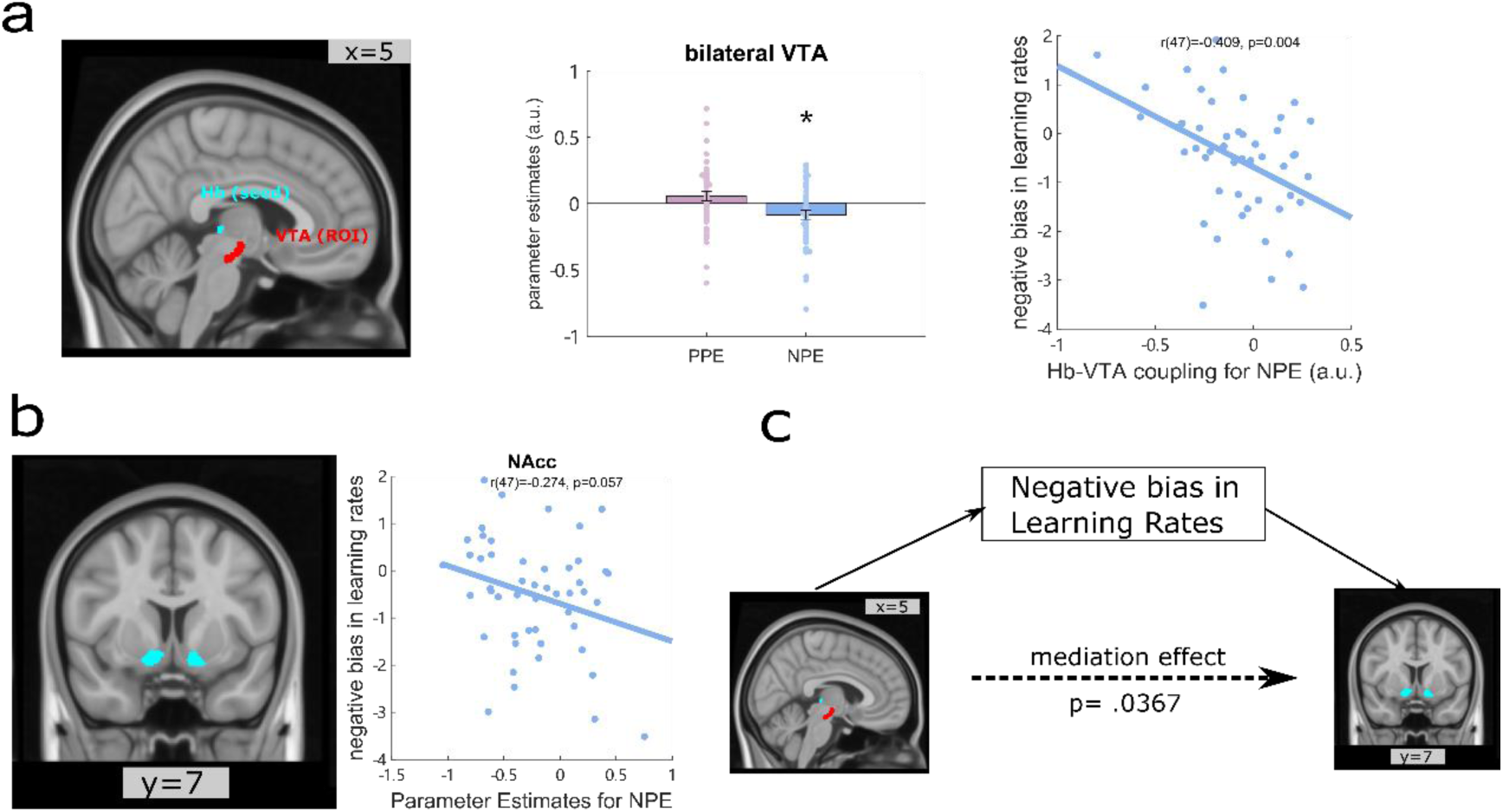
Psychophysiological interactions (PPI) result using bilateral habenula (Hb) as seed region. **a)** negative functional connectivity between the habenula and the VTA during negative prediction errors (NPE) correlated with negative learning bias. **Left:** a coronal view of the Hb and VTA anatomical mask. **Middle:** a bar graph depicting parameter estimates of Hb-VTA functional connectivity strength as a function of PPEs and NPEs, respectively. Each dot on the bar graph represents a parameter estimate for each participant. Error bars indicate standard errors (s.e.); *p < 0.05. Right: a scatter plot for the correlation between the Hb-VTA functional connectivity during NPEs encoding (x-axis) and negative bias in learning rates (y-axis) (p=.004). b) Left: a coronal view of the bilateral NAcc anatomical mask. Right: a scatter plot for the correlation between the parameter estimates of BOLD responses in the bilateral NAcc to NPE (x-axis) and negative bias in learning rates (y-axis) **c) mediation effect of learning rates bias.** Hb-VTA coupling impacted the NAcc BOLD responses to NPEs through the mediation of the learning bias (mediation p=.0367).

### Habenula-VTA functional connectivity and learning rates

Rodent evidence indicates that inactivating lateral Hb induces a positive learning bias, mediated via attenuation of an inhibitory response within VTA to reward omission, i.e. a negative prediction error (26). On this basis, we hypothesized that functional connectivity between Hb and VTA would relate to a learning rate bias. To test this, we ran a psychophysiological interaction (PPI) analysis, with bilateral Hb as the seed region (see Methods). This revealed a significant negative functional connectivity between Hb (seed) and bilateral VTA (ROI) during NPEs encoding (t(48)= -2.479, p=.017, see Fig. 3 a middle), consistent with evidence from animal studies (23, 26). Thus, the worse an outcome relative to expectation, the greater the negative coupling between Hb and VTA. Importantly, as predicted, we found a greater negative habenula-VTA connectivity during NPE encoding related to a more negative bias in learning rates (Fig3a right, r(47)=-.409, p=.004).

Given rodent evidence that Hb inhibits VTA (38, 39), our PPI results are consistent with enhanced inhibition from Hb to VTA in response to NPEs. To formally test this proposed mechanism, we used a structural equation model (SEM) to probe directionality of influence in connectivity between our a priori ROIs (Hb, VTA, NAcc and mPFC) (40, 41) (see supplementary method). Model evidence from SEM suggested that an interaction between ROI regions more likely originated from Hb than from any of the other three ROIs, with the winning model indicating the Hb influences VTA first, and NAcc and mPFC then both being subject to an influence from VTA (see Fig. S6). Finally, as we showed above an association between NAcc BOLD activation to NPEs and a learning rate bias, we probed this interaction in additional mediation analyses. This revealed a Hb-VTA functional coupling impacted the NAcc BOLD responses to NPEs through an influence on a learning bias (mediation p=.0367, see Fig. 3c).

### Variance enhances prediction error signals in the NAcc and the vmPFC

We hypothesized that a preference for broader options (PVB) emerges because higher variances boost a biased updating of option expected values (as suggested by our 2lr-RW model). On this basis, we tested whether prediction error signals were boosted for the broader compared to the narrower option. Thus, we ran a new first-level GLM design (see GLM 2 in the methods) on fMRI data, where we modelled prediction errors for the broader and narrower options respectively in a given block. Note that 1) this limited the analysis to equal mean blocks alone, where the two options were chosen approximately equally 2) we combined PPE and NPE as a single PE regressor for each option because of limited trials number for each. For prediction error signal for the chosen option, a variance (broad vs narrow) x mean (high vs low) repeated ANOVAs indicated a significant main effect of variance in bilateral NAcc (Fig 4 b, F(1,48)=15.298, p<.001) and vmPFC (Fig 4 e, F(1,48)=4.743, p=.034), but not in habenula or VTA (all p> .419). Additionally, we also ran the same analyses for the unchosen outcome which revealed that NAcc encoded prediction errors for the unchosen outcome in the opposite direction, i.e., lower the BOLD response to higher prediction errors (all t<-2.665, p<.010), where repeated ANOVAs also revealed a significant main effect of variance in bilateral NAcc (Fig 4 b, F(1,48)=4.302, p=.043), with a stronger variance negative modulation on BOLD signals for the broader option, an effect not seen for vmPFC (p=.250).

**Fig. 4.**
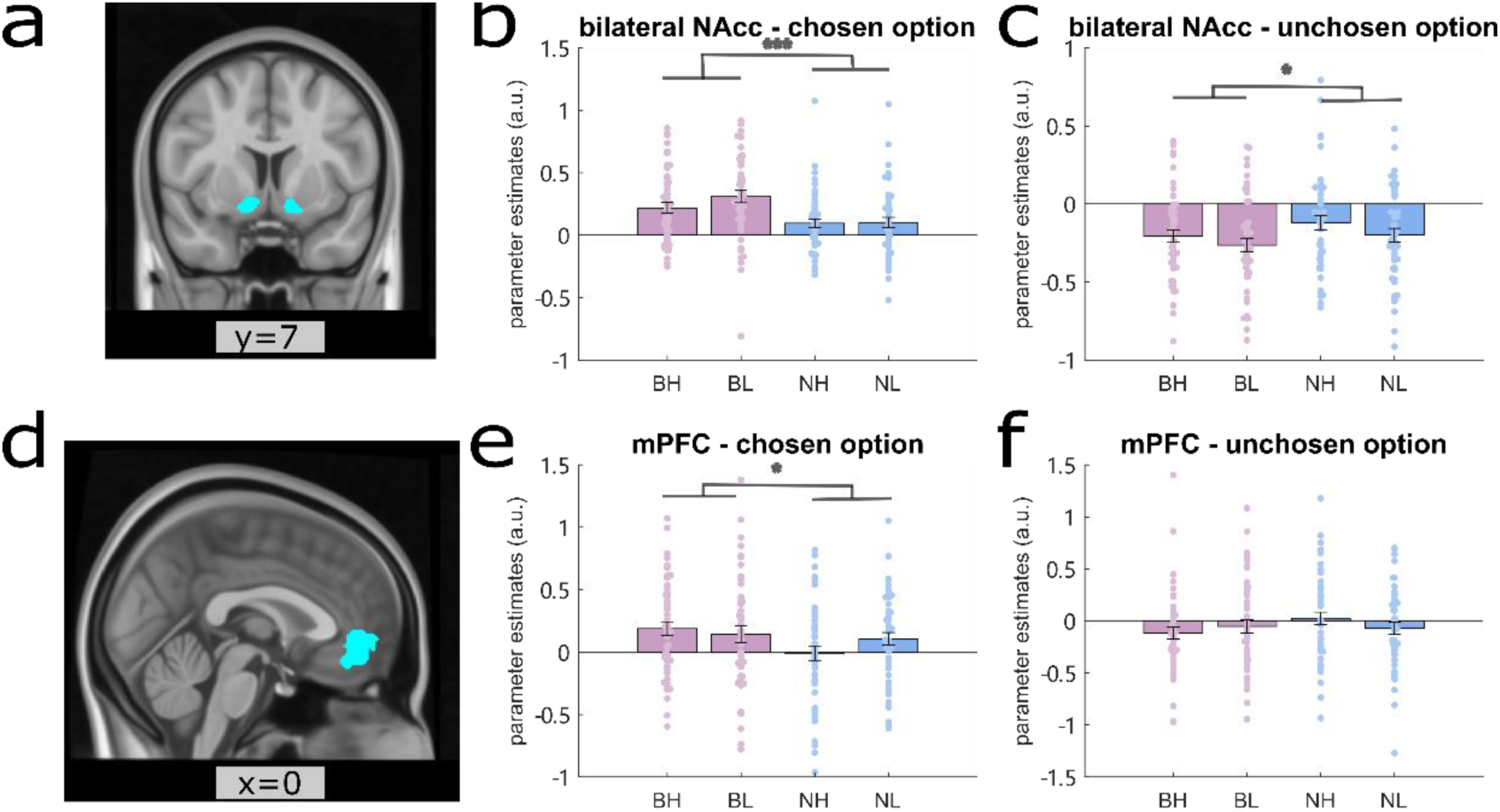
Variance boost prediction error signals in NAcc and vmPFC. **a)** a coronal view of the bilateral NAcc anatomical mask. **b-c)** parameter estimates of BOLD responses in this bilateral NAcc anatomical mask to prediction errors for each distribution type for the chosen (b) and unchosen option (c) respectively in the equal-mean blocks. **d)** a coronal view of the vmPFC mask. **e-f)** parameter estimates of BOLD responses in this vmPFC mask to prediction errors for each distribution type for the chosen (e) and unchosen option (f) respectively in the equal-mean blocks. BH: broad-high, BL: broad-low, NH: narrow-high, NL: narrow-low. Each dot on the bar graph represents a parameter estimate for each participant. Error bars indicate standard errors (s.e.); *p < 0.05; **p < 0.01; ***p < 0.001.

## Discussion

Combining a magnitude learning task and ultra-high resolution fMRI, we investigated mechanisms underlying the emergence and expression of individual differences in pro-variance bias. We show that individual differences in learning rate bias, derived from a 2lr-RW model, are associated with PVB, in turn associated with task-independent individual differences in anxiety and depression scores.

We identified regionally distinct responses to model-derived PPEs and NPEs in dopaminergic regions including NAcc, VTA and vmPFC, areas previously implicated in reward prediction error encoding in humans (see (42) for a meta-analysis). However, few studies have previously examined the joint expression of positive and negative PEs specifically. Notably, we show NAcc is positively modulated by PPEs and negatively by NPEs, a finding consistent with previous fMRI evidence showing ventral striatum sensitivity, including NAcc, to both unexpected reward and reward omission (43, 44). On the other hand, VTA was sensitive to PPEs but not NPEs, in line with previous human fMRI studies showing VTA expresses significant BOLD responses to unexpected rewards but not reward omission (44), and a PPE-like signal in an electric shock expectation task (44, 45). By contrast, we found vmPFC was significantly modulated by NPEs alone, consistent with other human-fMRI evidence that vmPFC BOLD responses were selective for unexpected reward omission (46) as well as the expression of an NPE signal in a fear conditioning task (47).

We show that human Hb encodes trial-by-trial NPEs, consistent with evidence that lateral habenula plays a key role in value-guided behaviour in non-human animals (for a review see (48)). Heretofore, its small size and relatively low resolution of 3T MRI have meant its role in human value-based learning has remained elusive. Studies using 3T fMRI report heightened BOLD responses in Hb to aversive outcomes, such as primitive negative events (shock) (49–52), negative feedback (53), punishment (54), a high probability of loss (55), and cued primary reward (juice) omission (56). However, studies have not shown Hb BOLD responses are modulated by monetary loss (49, 50), though a recent 7T fMRI study showed relative BOLD Hb deactivation to monetary loss avoidance (57). Here by combining ultra-high-resolution imaging and computational modelling, we go a step further and provide the first human evidence, to our knowledge, that Hb encodes trial-by-trail NPEs.

In our magnitude learning task, a negative bias in learning rates, as derived from a 2lr-RW model, captured individual differences in a pro-variance bias. This finding coupled with the observation that a more negative Hb-VTA functional connectivity during NPEs was associated with an enhanced negative bias in learning rates highlights the importance of this pathway in risk-aversive behaviour. Thus, we suggest that Hb-VTA modulation of a learning rate bias may contribute to the emergence of individual differences in risky behaviour. Evidence consistent with this mechanistic account includes evidence from rodents that stimulating a Hb-VTA pathway induces conditioned place aversion (58). Note that the resolution of human non-invasive neuroimaging does not enable examination of the lateral habenula specifically, and our region of interest includes both lateral and medial habenula. Lateral habenula alone is linked to expression of NPEs, while medial habenula has a different functional attribution (59–61). However, despite this limitation in resolution, we provide evidence that human habenula encodes NPEs and through its functional connectivity to VTA exerts an influence on negative bias learning.

An enhanced negative BOLD response in NAcc to NPEs was associated with a more negative learning rate bias. Consistent with our results, we note a previous fMRI study showed that a differential NAcc BOLD response to prediction errors for risky options, relative to certain options, is linked to behaviour risk preferences (62). In the aversive domain, a previous finding from our lab also showed a negative modulation of striatal responses by aversive prediction errors (delivery of shock) linked to a more negative learning bias and a reduced propensity to gamble (63). At a mechanistic level, our mediation analysis suggests that Hb-VTA coupling impacts NAcc responsivity to NPEs via an influence on learning rates. We speculate that the association between a striatal involvement and a learning bias, as seen both here and in an earlier study (63), may reflect a modulation by upstream reward regions, specifically a functional interaction between Hb and VTA.

Intriguingly, we found evidence for adaptive coding of reward prediction errors in NAcc and vmPFC. Specifically, in both regions, item-based reward prediction error signals were enhanced under higher variance, consistent with an uncertainty boosting of RPE signals. Consistent with our results, uncertainty has been shown to increase an outcome signal in NAcc (64). In non-human primates, both reward risk (variance) and reward volatility (unexpected uncertainty) increase neural activity in the orbital frontal cortex (OFC) (65, 66), a region with functional attributions similar to vmPFC in humans (67). However, other evidence, using an instructed cue learning task whereby participants were informed regarding the standard deviation (SD) of outcomes, showed this served to reduce RPE signals in the ventral striatum (68). We suggest that, as in our task, when subjects are actively learning variances then this serves to boost prediction error signals, as opposed to when variances are well-learned or instructed, which results in a consequential decrease in prediction error signals. Indeed, a study compared learnt and described risk during a risk decision-making task showed that the expected value signal is higher in vmPFC for the learnt risky option (69). Consistent with this is evidence that for a positive encoding of uncertainty during exploration (in learning) and negative encoding during exploitation (well learnt) in the vmPFC (70). Interestingly, we showed that the RPE for the unchosen option is also modulated by variance, however, this effect is not seen for vmPFC.

A reduced pro-variance bias, and a negative bias in learning rates, were associated with higher anxiety and depression scores. Negative cognitive bias has been a core cognitive model of depression (71). We show that negative learning bias is modulated by Hb-VTA functional coupling to NPEs and this aligns with evidence indicating Hb plays a major role in depression (72). For example, an animal depression model (cLH rats) has shown increased spontaneous lateral habenula activity (bursting), which can be attenuated by ketamine, a fast-acting antidepressant (73). Notably, in macaque monkeys, ketamine has been shown to increase pro-variance bias (11) while in rodents inactivation of Hb leads to more positive bias learning accompanied by reduced de-activities to reward omission in DA neurons in VTA (26). Together, these results showed a link between the habenula-dopamine pathway to negative bias in depression.

In conclusion, we show that a negative bias in learning rates, which accounts for individual differences in pro-variance bias, relates to an enhanced Hb-VTA functional coupling for NPEs. By modulating learning rate bias, the Hb-VTA functional interaction influences downstream region NAcc’s response to NPEs. The results link learning signals in the brain and PVB and provide a neural basis for individual differences in risk preference, with implications for psychopathology.

## Methods

### Participants and procedures

In an fMRI study, we recruited 57 participants from the general population. To cover a spectrum of anxiety and depression levels, participants were prescreened on their scores on anxiety and depression questionnaire scores, measured by the Zung self-rating depression scale (SDS) and the state-trait anxiety inventory (STAI). We adopted the same behaviour data control criteria as our previous study (8). More specifically, five subjects were excluded from the analyses because their mean accuracies for the different-means blocks were less than 60%; one was excluded because they chose the same option for all the trials in at least one of the four same-mean blocks; a final subject was excluded for both reasons. One additional participant was excluded on the basis that they showed a pro-variance bias that exceeded 3 standard deviations of the overall group mean. This resulted in a final sample of 49 (34 females) participants, aged 18-30.

The experiment was implemented using the software PsychoPy (v2021.1.4) (74). Each participant completed 8 blocks of the learning task in a 7T MRI scanner, with a rest between blocks. Participants were reimbursed a base amount plus a bonus earned during the learning task. All participants completed 5 questionnaires on the scan day measuring different cognitive traits associated with anxiety and depression (75): 1) the state-trait anxiety inventory (STAI) (76); 2) the Zung self-rating depression scale (SDS) (77); 3) the rumination response scale (RRS) (78); 4) the intolerance of uncertainty scale (IUS) (79); 5) the 17-item Dysfunctional Attitude Scale form A (DAS-A) (80). We used their questionnaire scores taken on the scanning day for all the relevant analyses.

All participants provided informed consent before any testing. This study was approved by the Beijing Normal University Research Ethics Committee (IRB_A_0051_2021001) in accordance with the Declaration of Helsinki(81).

### The experiment task

We adapted a magnitude learning task, as reported in a previous paper (8), wherein participants made choices between two card decks with varying hidden card number distributions for 30 trials in a block (Fig. 1 b). For each block, the same two card decks, cued by different colours and/or patterns of the back of the poker cards, were shown throughout. Two new card decks (different colours and/or patterns from the ones shown before) were assigned to each new block. There were 8 blocks in total: 4 different-mean blocks: one narrow-high vs broad-low (NHBL) block, one broad-high vs narrow-low (BHNL) block, and two broad-high vs broad-low (BHBL) blocks; and 4 same-mean blocks: two broad-high vs narrow-high (BHNH) and two broad-low vs narrow-low (BLNL). Note that, compared to the previous paper, we did not include the bimodal block which didn’t show any correlation with individual differences in anxiety-and-depression-related traits (see (8) for more details). The block order was counterbalanced across participants.

At the beginning of each trial, participants were presented with two pictures of the back of the card decks and asked to choose (self-paced) one of them. The positions (left or right side) of the two card decks were randomized from trial to trial. When they completed a choice, the cross in the centre of the screen changed into a triangle and pointed to the card deck they had chosen for the trial. One to three (mean 2s) seconds after a choice, the card number from one of the decks was revealed for 1-3s (mean 2s) and then turned back. Half a second later, the card number from the other card deck was revealed for 1-3s (mean 2s). The order was counter-balanced across trials such that half of the time one card deck was revealed first. Finally, the card numbers were shown together again for 1s. Simultaneously, participants were shown how many points they had won or lost for that trial. If the chosen card number were higher than the unchosen card number, participants gained points equal to the value difference between the two numbers for that trial; if the chosen card number was lower than the unchosen one, participants lost points equal to the value differences between the two cards. If the two card numbers were equal the total points did not change for that trial. To encourage better performance, participants were told that a bonus would be added to their final reimbursement proportional to the points they had won by the end of the experiment.

### Behavior data analysis

#### Statistical analysis

The percentages of choosing the higher option (for the different-mean blocks) or the broader option (for the same-mean blocks) were calculated for each block type respectively. The mean pro-variance biases were calculated by averaging the percentages of choosing the broader options for all 4 same-mean blocks, i.e., two BHNH blocks and two BLNL blocks. Using one-sample T-tests, we tested whether participants’ percentages of choosing the higher or broader option were different than the chance level (50%) for each block type, respectively. Paired T-tests were used to compare participants’ overall performance between different block types. Correlation analyses were performed using Pearson’s Correlation. The statistical analyses were performed in IBM SPSS Statistics, version 26.0.0.0®.

#### Computational modelling

Our winning model is a two learning rates Rescorla–Wagner model (2lr-RW) (82), a simple variant of the RW model (12). In this model, two different learning rates (α_+_ and α_-_) are assigned to the positive and negative prediction errors respectively (Equation 1). More specifically, for every trial, using Equation 2, a prediction error (δ) was calculated using the outcome received for the current trial (R_t_) minus the expected value for the current trials (V_t_). The expected value for the next trial (V_t+1_) is updated using Equation 3 by taking the EV from the previous trial (V_t_), adding the product of prediction error times and a learning rate α(δ) dependent on the sign of the prediction error for the current trial: α_+_ is used if this prediction error is positive (i.e. higher than or equal to 0); α_-_ is used if this prediction error is negative (i.e. lower than 0),(see Equation 1).

Bias in learning rates would lead to a bias in updating distribution variances, making the expectation (V_t_) higher or lower than the true mean of a distribution. The negative learning rate bias (LR_neg-bias_) is calculated using **Equation 4**. In previous work, we showed in simulation the learning rate bias would generate a spectrum of pro-variance biases in the paradigm deployed in the current study (8). The outcome values (i.e. 1-13) were rescaled to 0.01-0.99 and the prior expected value was set to 0.5.

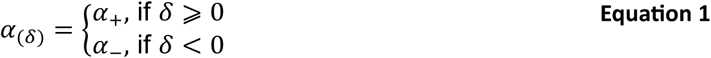

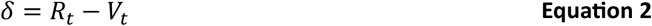

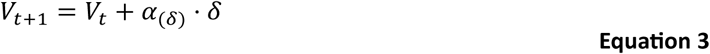

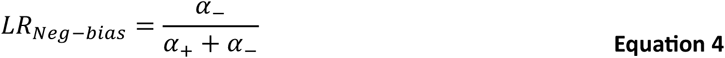

For a two-arm bandit choice task, the probabilities of choosing an option are linked to the difference between the learned expected values for the two options. We adopted a standard softmax function for decision-making processes to generate a probability of choosing option a (Equation 5) (83) where nverse temperature β controls the stochasticity of decision-making. Note that Equation 5 is also used in other models in the current study (see Supplementary for details of other models used).

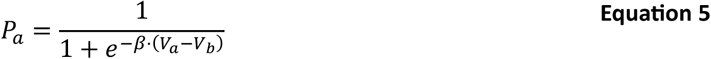

All models were fitted in Matlab R2020b using a variational Bayes approach. Only behavioural data from the four same-mean blocks were used for the model fitting. This is because the different-mean blocks were relatively simple, and a majority of participants always chose the higher option in these blocks. All trials from the same-mean blocks were used in modelling fitting. Bayesian information criterion (BIC) was calculated for each model using the best-fitted parameters for each participant (Equation 6 & Equation 7). L^^^ denotes the maximized value of the likelihood function of the model M, x: the observed data, n: the number of observations, k: the number of free parameters in the model. BICs for a random model were calculated with each decision probability set as 0.5 and k=0. BIC scores were summed across participants, with lower sum BIC indicating better model fit. Delta BIC for each model was calculated by subtracting the BIC score of the best-fitting model in each experiment.

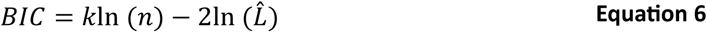

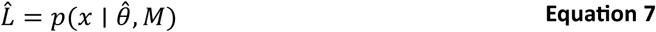

We then used the best fitted parameters for each participant to calculate the trial-by-trial prediction errors that used in GLM 1 for the first-level fMRI data analyses.

### fMRI data analysis

#### Imaging data acquisition

Imaging data was collected using a Siemens 7T MRI scanner in Peking University in Beijing, China. Functional data were acquired using a multiband gradient-echo T2* echo planar imaging (EPI) sequence with a 1.2*1.2*1.2 mm resolution; multiband acceleration factor 2; repetition time (TR) 1375 ms; echo time (TE) 19 ms; flip angle 60°; and a GRAPPA acceleration factor 3. Field of view (FOV) was adjusted to cover the basal ganglia and thalamus, see Fig S3. During the main functional runs, cardiac and respiratory frequencies were collected to regress out the effect of physiological noise. Additionally, a single whole-brain functional image was acquired after the main functional runs with the same parameters except for a TR of 3298 ms.

Structural data were acquired with a T1-weighted MP-RAGE sequence with a 0.7*0.7*0.7 mm resolution; GRAPPA acceleration factor 2; TR 2200ms; TE 1.65 ms; and inversion time (TI) 1050ms. A separate Fieldmap sequence was acquired with 2*2*2 mm resolution; TR 620 ms; TE1 4.08 ms; TE2 5.10 ms.

#### fMRI data preprocessing

Pre-processing was performed using tools from FMRIB Software Library (FSL) (84). Functional images were first normalized, spatially smoothed (Gaussian kernel with 2 mm full-width half-maximum (FWHM)), and temporally high-pass filtered (3 dB cut-off of 90s). Participants’ head motion during the scanning was removed by MCFLIRT (85). The Brain Extraction Tool (BET) (86) was used on functional and structural images to separate brain from non-Brian matter. Structural images were bias field corrected before BET. The functional images were registered to Montreal Neurological Institute (MNI)-space in three stages: 1) partial FOV task EPI to whole-brain EPI using FMRIB’s Linear Image Registration Tool (87). 2) Whole-brain EPI to individual structural image using Boundary-Based Registration (BBR) (88) by incorporating Fieldmap correction. 2) Individual structural image to Standard image by using FMRIB’s Non-Linear Image Registration Tool (FNIRT).

#### Model-based fMRI data analyses

Statistical analyses of the fMRI data were performed at three-levels using FSL FEAT (89). At the first level, univariate general linear models (GLMs) were run for each session/block for each participant.

#### GLM 1

BOLD=β_0_+ β_1_*Response at choice phase + β_2_*reaction time at choice phase+ β_3_*mean of choice phase + β_4_* positive prediction errors for the chosen option + β_5_* mean of the trials for the chosen option with positive prediction errors+ β_6_* negative prediction errors for the chosen option + β_7_* mean of the trials for the chosen option with negative prediction errors + β_8_* positive prediction errors for the unchosen option + β_9_* mean of the trials for the unchosen option with positive prediction errors+ β_10_* negative prediction errors for the unchosen option + β_11_* mean of the trials for the unchosen option with negative prediction errors + β_12_*mean of the 3^rd^ feedback. (note that negative prediction errors were flipped in signs so that the higher the number the more the outcome negatively deviated from the expected value.)

#### GLM 2

BOLD= β_0_+ β_1_*Response at choice phase + β_2_*expected value difference between the two options at choice phase + β_3_*mean of choice phase + β_4_* prediction errors for the broader option when it was chosen and its outcome revealed + β_5_* mean of the trials for the broader option when it was chosen and its outcome revealed + β_6_* prediction errors for the broader option when it was not chosen and when its outcome revealed + β_7_* mean of the trials for the broader option when it was not chosen and when its outcome revealed + β_8_* prediction errors for the narrower option when it was chosen and when its outcome revealed + β_9_* mean of the trials for the narrower option when it was chosen and when its outcome revealed + β_10_* prediction errors for the narrower option when it was not chosen and when its outcome revealed + β_11_* mean of the trials for the narrower option when it was not chosen and when its outcome revealed + + β_12_*mean of the 3^rd^ feedback.

For GLM 1, at the second level, the contrast of parameter estimates from all sessions/blocks was combined for each participant using fixed effects. For GLM 2, at the second level, the contrast of parameter estimates from the same block type (i.e. the two BHNH blocks and two BLNL blocks) was combined for each participant using fixed effects.

The subject means from the second level were put into a mixed effects model (FLAME1) (90) for the group-level analyses.

#### Region of interests

The habenula ROIs were manually drawn on each participant’s high-resolution structural scans according to an established protocol (37). The Hb ROIs in individual spaces were then transferred to standard MNI space and combined to create a study-specific Hb ROIs for left, right and bilateral Hb ROI masks respectively. The Hb ROIs in the MNI space were thresholded such that the voxels were shared by more than 30% of the participants (i.e. 15). Additionally, we used anatomical masks [28] for NAcc and VTA, and a functionally defined ROI for the ventral medial prefrontal cortex (vmPFC) based on a meta-analysis of value-based decision-making tasks (see Fig. 2 b-d) (36).

#### Psychophysiological interaction analysis

In order to examine task-related interregional interactions among brain regions, that were modulated by PPEs and NPEs, we implemented psychophysiological interaction (PPI) analyses using FSL (91) and where bilateral Hb ROI was the seed region. Based on **GLM 1**, we ran PPI for the NPEs and PPEs parametric regressors respectively.

#### Mediation analysis

A mediation analysis using the multilevel mediation and moderation (M3) toolbox (92) in Matlab with permutations (n=50000) was performed to examine the relationships amongst Hb-VTA functional connectivity, NAcc BOLD responses to NPE and a negative bias in learning rates. In the analysis, negative biases in learning rates were set as the mediation factor, and Hb-VTA PPI connectivity to NPEs and NAcc BOLD modulation parameter estimates by NPEs as independent and dependent variables respectively.

## Data Availability Statement

Anonymized behavioral and ROI data have been deposited in the Open Science Framework (DOI 10.17605/OSF.IO/2KCXN).

## Acknowledgements

We thank Dr. Leo Chi U Seak for providing valuable feedback on manuscript drafts.

## Authors’ contributions

WL and RD designed research; WL, XZ and JX performed research; WL and JX analyzed data; and WL, RD and JX wrote the paper.

