## Supplement Materials for "Habenula-ventral tegmental area functional coupling and risk-aversion in humans"

### Supplementary Results

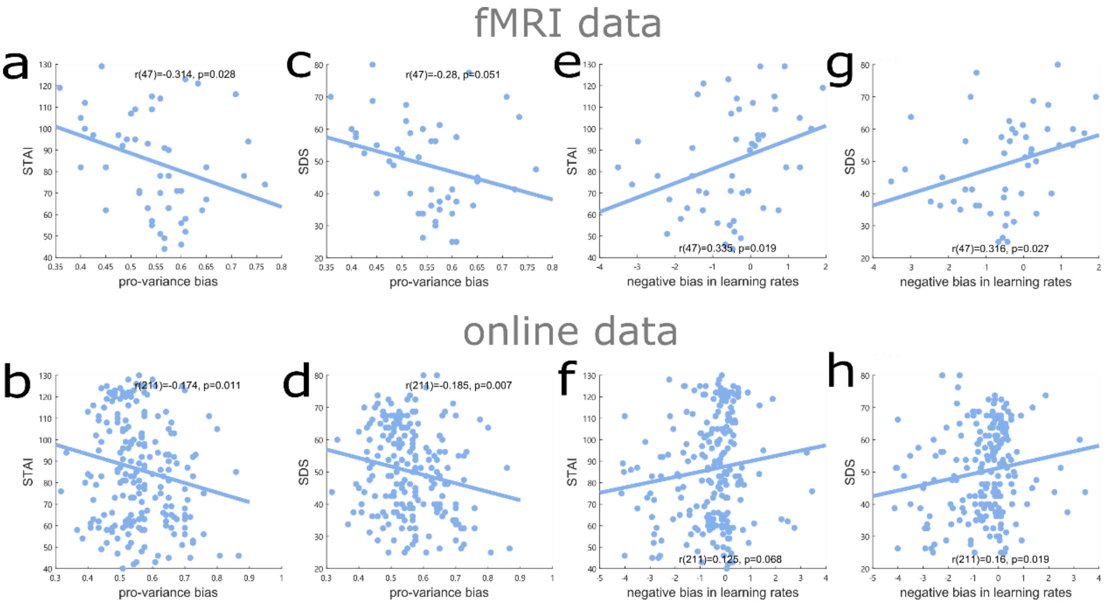


**Fig.S 1 | correlations with questionnaires. a-b)** PVB negatively correlated with State-Trait Anxiety Inventory (STAI) scores for the fMRI data (a) and online data (b). **c-d)** PVB negatively correlated with the Zung Depression Scale (SDS) scores for the fMRI data (c) and online data (d). **e-f)** Negative bias in learning rates correlated with State-Trait Anxiety Inventory (STAI) scores for the fMRI data (e) and online data (f). **g-h)** Negative bias in learning rates correlated with the Zung Depression Scale (SDS) scores for the fMRI data (g) and online data (h).

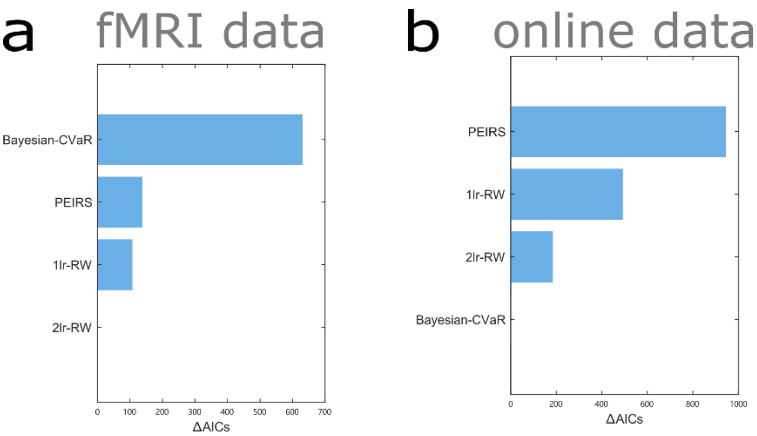


**Fig.S 2 | Model comparisons.** The relative Akaike information criterion (AIC) results for the fMRI data (a) and online pilot data (b).

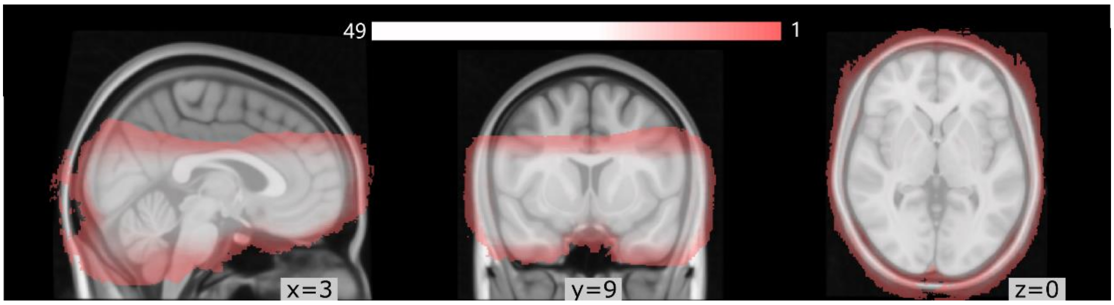


**Fig.S 3 | Field-of-view (FOV) for each participant after registration overlayed on a standard brain.** Colormap indicates the number of participants.

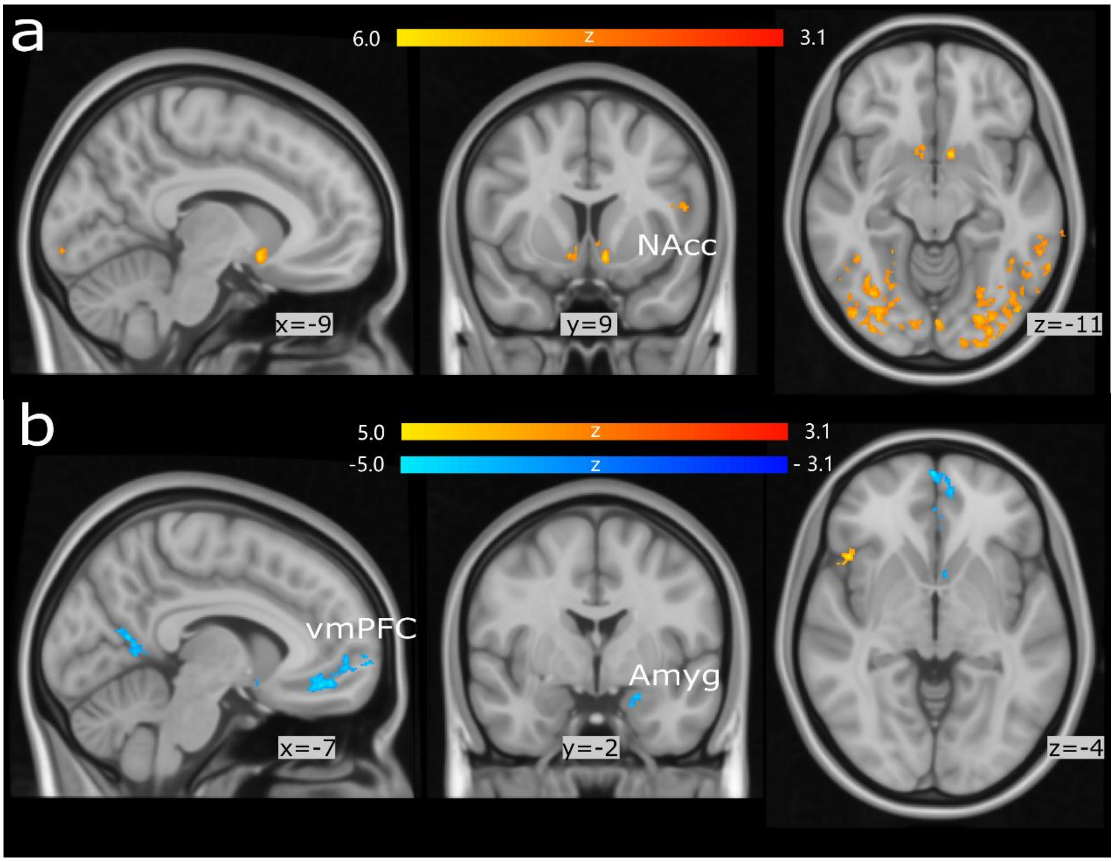


**Fig.S 4 | Whole FOV results for trial-by-trial PPEs (a) and NPEs (b) parametric regressors.** Cluster p<0.05, |Z|>3.1. NAcc: nucleus accumbens; vmPFC: ventral medial prefrontal cortex; Amyg: Amygdala.

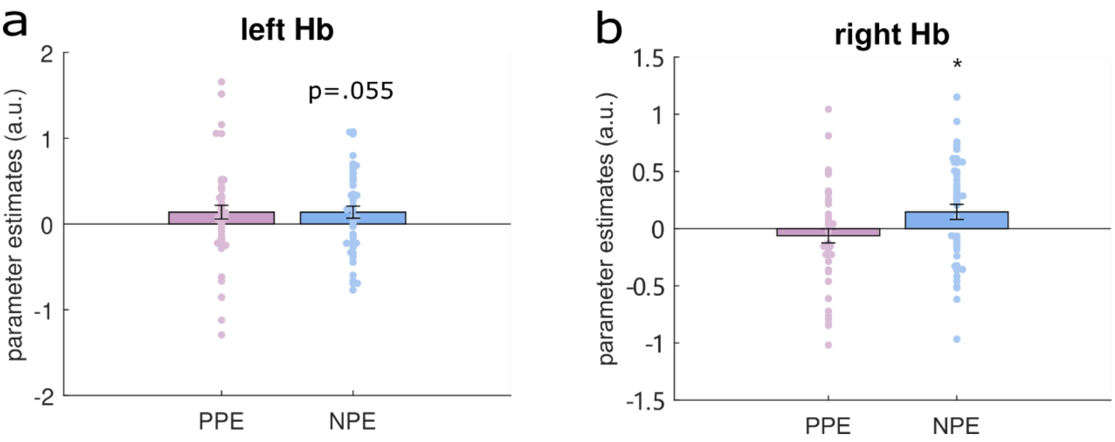


**Fig.S 5 | Left and right habenula (Hb) responses to PPE and NPE.** BOLD responses in both the left (a)

and right (b) Hb were positively modulated by negative prediction errors (NPEs) but not positive prediction errors (PPEs).

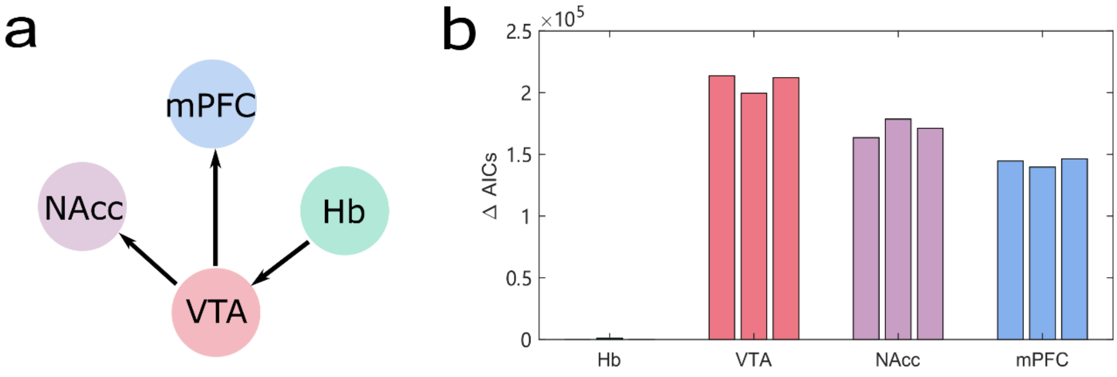


**Fig.S 6 |Structural Equation Modeling (SEM) results. a)** the winning model. **b)** the model comparison (AICs) results for models starting from one of the four ROIs. The x-axis label denotes the starting ROI region for each model.

#### Model free fMRI results show similar results as the model-based fMRI analyses

In this magnitude learning task design, participants win or lose a different number of points from trial to trial, which enables us to investigate parameter modulation of the outcomes without using computational model estimates. Therefore, we examined the BOLD responses to win amount and loss amount (GLM3, see supplementary methods). As expected, BOLD responses in the NAcc (**Fig.S 7**) increased as participants won more points in a trial (i.e. win amount, t(48)=6.018 p<.001) and de-activated more for losing more points (i.e. loss amount, t(48)=-4.405 p<.001). Similar to the model-based fMRI results, the habenula showed marginally significant positive modulation by loss amount in the bilateral habenula (t(48)=1.849 p=.071, left habenula (**Fig.S 7**): t(48)=2.158, p=.036, right habenula： p=.231). However, mPFC (**Fig.S 7**) showed significantly positive modulation by win amount (t(48)=2.724 p=.009) and negative modulation by loss amount (t(48)=-2.106 p=.041). The whole-FOV analysis showed similar results as the ROI analysis (see **Fig.S 8**).

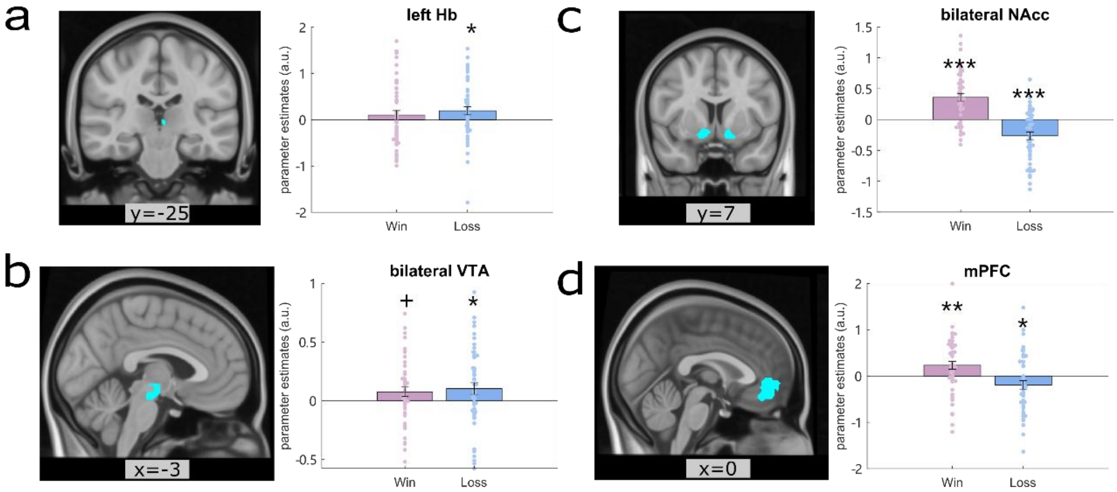


**Fig.S 7 | regions respond to model-free outcome magnitudes.** **a)** BOLD responses in the left habenula (Hb) are positively modulated by loss amount (p=.036) but not by win amount (p=.785). Left: a coronal view of the left Hb anatomical mask. Right: parameter estimates of BOLD responses to win and loss amount respectively in this left Hb anatomical mask. **b)** BOLD responses in the bilateral ventral tegmental area (VTA) showed a positive modulation by win amount (p=.065) and loss amount (p=.040). Left: a sagittal view of VTA anatomical mask. Right: parameter estimates of BOLD responses to win and loss amount respectively in this bilateral VTA anatomical mask. **c)** BOLD responses in the bilateral nucleus accumbens (NAcc) positively modulated by win amount and negatively by loss amount (both p<.001). Left: a coronal view of the bilateral NAcc anatomical mask. Right: parameter estimates of BOLD responses to win and loss amount respectively in this bilateral NAcc anatomical mask. **d)** BOLD responses in the medial prefrontal cortex (mPFC) are positively modulated by win amount (p=.009) and negatively by loss amount (p=.041). Left: a sagittal view of the mPFC functional defined mask. Right: parameter estimates of BOLD responses to win and loss amount respectively in this mPFC mask. Each dot on the bar graph represents a parameter estimate for each participant. Error bars indicate standard errors (s.e.); + p<.1 *p < 0.05; **p < 0.01; ***p < 0.001.

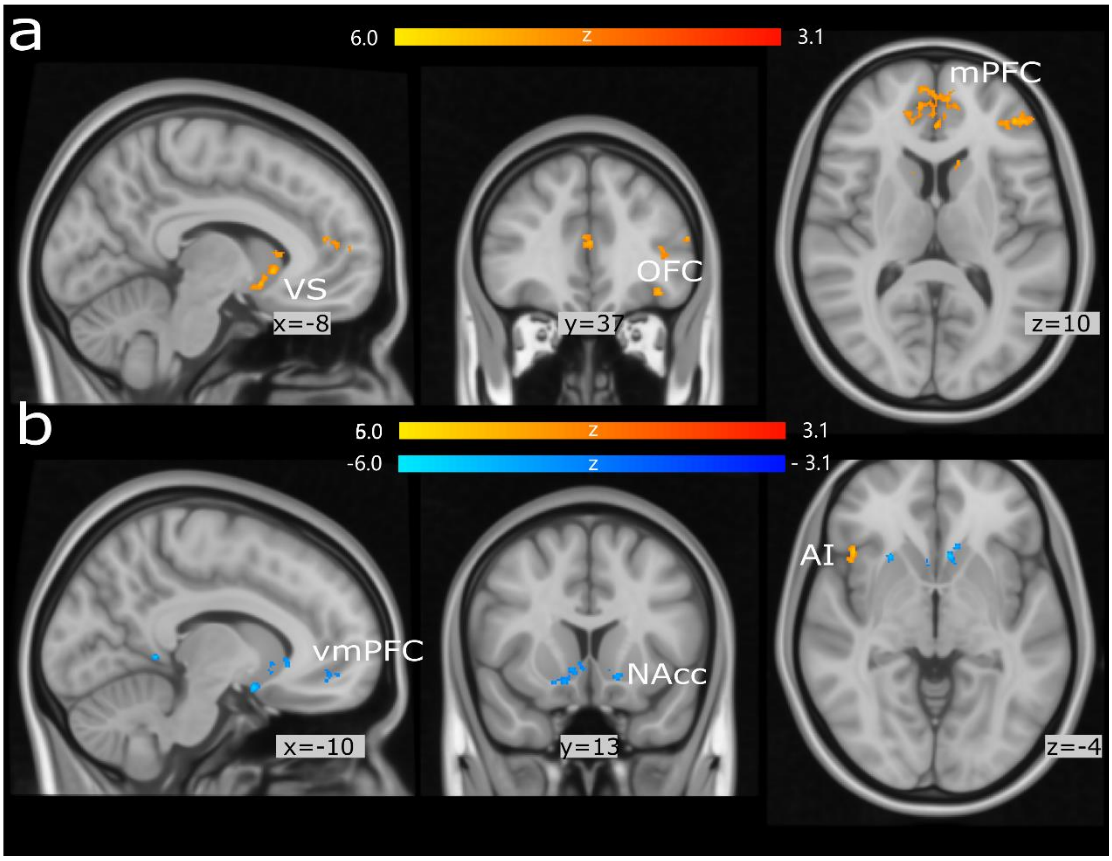


**Fig.S 8 | Whole FOV results for trial-by-trial win (a) and loss (b) amount parametric regressors.** Cluster $p<0.05$, $|Z|>3.1$. VS: ventral striatum; OFC: orbital frontal cortex; mPFC: medial prefrontal cortex; vmPFC: ventral medial prefrontal cortex; NAcc: nucleus accumbens; AI: anterior insula.

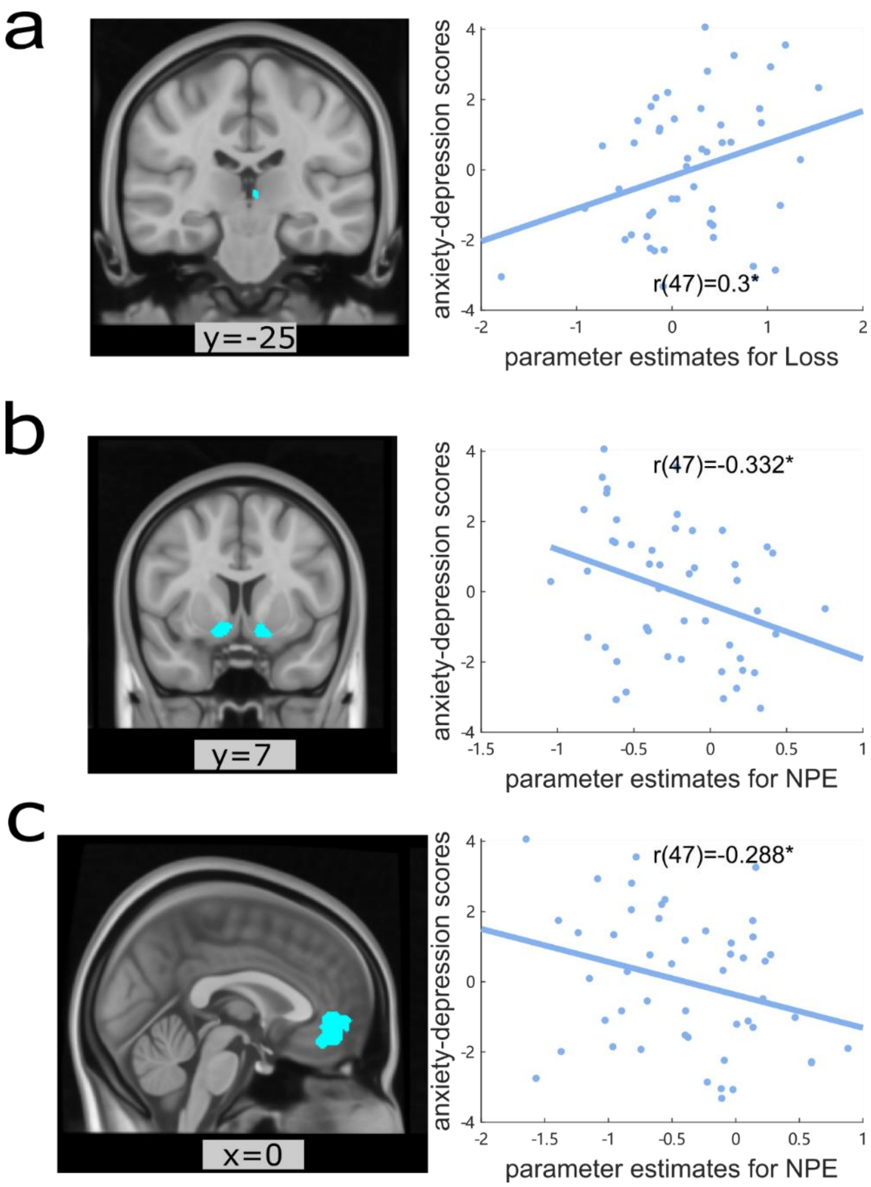


**Fig.S 9 | Neural correlates of negative events with anxiety-depression scores.** **a)** BOLD responses to loss amount in the left habenula (Hb) positively correlate with the anxiety-depression scores (p=.036). Left: a coronal view of the left Hb anatomical mask. Right: a scatter plot for parameter estimates of BOLD responses in the left Hb to loss amount (x-axis) and anxiety-depression scores (y-axis). **b)** BOLD responses to negative prediction errors (NPEs) in the bilateral nucleus accumbens (NAcc) negatively correlate with the anxiety-depression scores (p=.020). Left: a coronal view of the bilateral NAcc anatomical mask. Right: a scatter plot for parameter estimates of BOLD responses in the NAcc to NPEs (x-axis) and anxiety-depression scores (y-axis). **c)** BOLD responses to negative prediction errors (NPEs) in the medial prefrontal cortex (mPFC) negatively correlate with the anxiety-depression scores (p=.045). Left: a coronal view of the mPFC functional mask. Right: a scatter plot for parameter estimates of BOLD responses in mPFC to NPEs (x-axis) and anxiety-depression scores (y-axis).

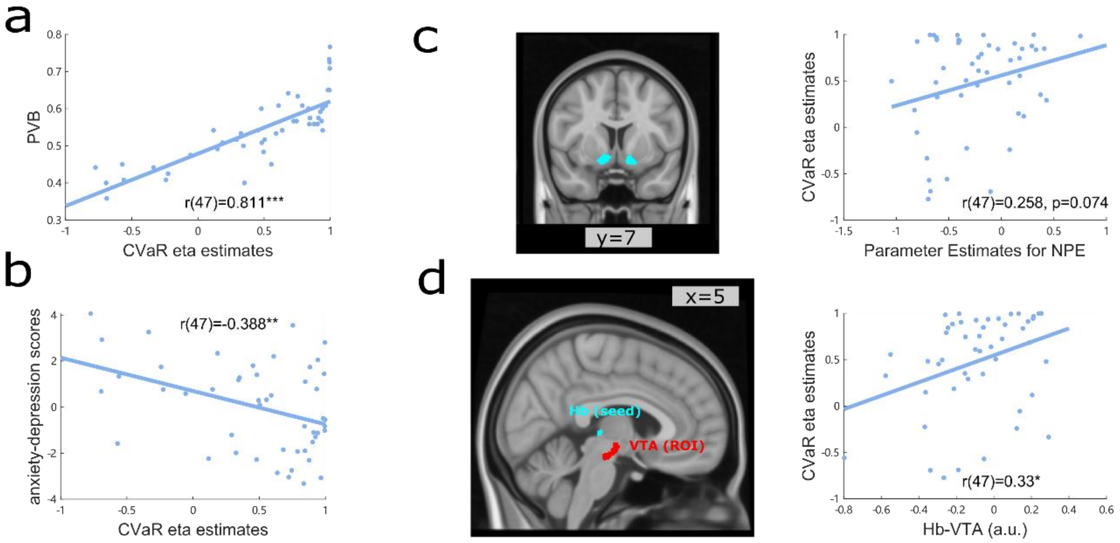


**Fig.S 10 | Correlations results for the eta estimated from the Bayesian-CVaR model.**

#### Variances represented in the mPFC

A recent rodent study using neuropixels data showed an abstract encoding of variance in NAcc, manifesting in a higher representational similarity between options that have similar distributions, i.e., higher within-distribution similarity than across-distribution [1]. Using representational similarity analysis (RSA) [2] we tested for such distributional encoding in our human fMRI data. To regress out an effect of variance in response to prediction errors (parametric modulation regressors, shown in Fig 4 b&e), we ran RSA on the contrasts for the mean regressors for the chosen option for each distribution type for the same-mean blocks and for each ROI respectively. Within both NAcc and vmPFC ROIs, we found evidence of increased representational similarity between broad-high (BH) and narrow-high (NH), as well as between broad-low (BL) and narrow-low (NL) (all $t(48)>12.89$, $p<.0001$). Additionally, in mPFC we found significantly high similarities between both broader options (broad-high (BH) and broad-low (BL)) (see **Fig.S 11** d, $t(48)=2.900$, Bonferroni adjusted $p=.035$). This pattern of findings is consistent with a suggestion that mPFC encodes a distribution of option outcomes, as suggested by distributional reinforcement learning theory [3].

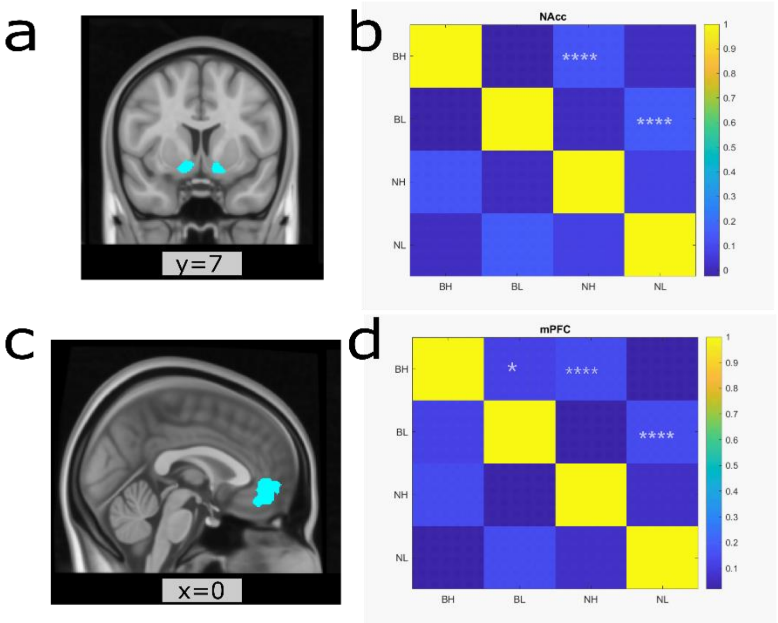


**Fig.S 11 | Neural representations of variances. a)** a coronal view of the bilateral NAcc anatomical mask. **b)** representation similarities in this NAcc mask between each type of option (option mean regressor) in the equal-mean blocks. **c)** a coronal view of the vmPFC mask. **d)** representation similarities in this vmPFC mask between each type of option (option mean regressor) in the equal-mean blocks. Each dot on the bar graph represents a parameter estimate for each participant. Error bars indicate standard errors (s.e.); *p < 0.05; **p < 0.01; ***p < 0.001;****p < 0.0001.

### Supplementary Discussion

We showed that in this fMRI sample the best fitted model was the 2lr-RW model. In previous paper and the pilot online study in China, all three independent samples we found that the Bayesian-CVaR model was the best fitting model. This might be a results of removing the bimodal blocks in the fMRI study or relatively reduced participants sized in the fMRI study compared to all other three samples. Nevertheless, we showed in Fig.S 10 that all the main results we shown for the learning rate bias estimates from the 2lr-RW model, also held true for the CVaR eta estimates from the Bayesian-CVaR model buy weaker.

### Supplementary Methods

#### Online pilot study

Before we conducted the fMRI study, we ran an online pilot study in China on a Chinese online study platform https://www.naodao.com/. The online pilot study used the same experiment design as described in a previous paper [4]. Part of this pilot study was also included in the supplementary of the paper [4]. The experiment was implemented using the software PsychoPy (v2021.1.4) [5]. In total, 320 participants were recruited for the online study. 107 participants were excluded based on the same pre-set criteria used here and in the previous paper [4].

#### Other computational models

**1lr-RW.** In this model, there is only one free parameter during the learning process, i.e., α, which controls how much the prediction errors (δ) are updated into the expected values (see Equation 1). Prediction errors are the differences between the outcome and the expected value for a given option.

$$V_{t+1} = V_t + \alpha \cdot \delta \quad \text{Equation 1}$$

$$\delta = R_t - V_t \quad \text{Equation 2}$$

**PEIRS.** In the PEIRS model, the expected values V and expected spread S of the outcomes are learned simultaneously using Equation 3 and Equation 4, respectively. The estimated spread is combined with the overall prediction errors of the options $\delta_{option}$ to determine the value V' (see Equation 6) used for the decision process (Equation 7). $S^t_{option}$ is the overall prediction of mean values of the two options offered compared to a global mean, which is 0.5 in this task (Equation 5). For example, for the both-high blocks, $S^t_{option}$ would be overall positive, $S^t_{option}$ for the both-low blocks, would be overall negative. ω in the Equation 6 controls the direction and how much the estimated spread would influence the decision-making process.

$$V_{t+1} = V_t + \alpha_Q \cdot \delta_{outcome} \quad \text{Equation 3}$$

$$S_{t+1} = S_t + \alpha_s \cdot (|\delta_{outcome}| - S_t) \quad \text{Equation 4}$$

$$S^t_{\text{options}} = \frac{V^t_a + V^t_b}{2} - 0.5 \quad \text{Equation 5}$$

$$V'_t = V_t + tanh(\omega \cdot \delta^t_{options}) \cdot S_t \quad \text{Equation 6}$$

$$P_a = \frac{1}{1 + e^{-\beta \cdot (V_a' - V_b')}}$$ Equation 7

**Bayesian-CVaR.** In this model, a probability density function θ of the value distribution for each option is learned using Bayes' rules. The posterior belief $P(\theta_{t+1})$ of the value distribution is updated trial by trial by the combination of the prior belief $P(\theta_t)$ and probability density function of the evidence for that trial $P(R_t)$ using Equation 8. The initial belief $P(\theta_0)$ is set as a flat distribution using a Beta distribution Beta (1,1). The probability density function of the evidence is a Beta distribution Beta (event$\alpha_t$, event$\beta_t$) with a mean of $R_t$ and the same variance (denoted as updatevar), as shown in Equation 10 & Equation 11, respectively. event$\alpha_t$ and event$\beta_t$ can be calculated using Equation 12 & Equation 13 respectively (these two equations were derived using the solve equation function on Equation 10 and Equation 11 in MATLAB).

$$P(\theta_{t+1}) \propto P(R_t) \cdot P(\theta_t)$$ **Equation 8**

$$P(R_t) = Beta(event\alpha_t, event\beta_t)$$ **Equation 9**

$$R_t = \frac{event\alpha_t}{event\alpha_t + event\beta_t}$$ **Equation 10**

$$updatevar = \frac{\mathrm{event}\alpha_t \cdot \mathrm{event}\ \beta_t}{(\ \mathrm{event}\alpha_t + \ \mathrm{event}\beta_t)^2 \cdot (\mathrm{event}\alpha_t + \mathrm{event}\beta_t + 1)}$$ **Equation 11**

$$event\alpha_t = -R_t \cdot (R_t^2 - R_t + updatevar)./updatevar$$ **Equation 12**

$$event\beta_t = (R_t - updatevar + R_t \cdot updatevar + 2 \cdot R_t^2 + R_t^3)./updatevar$$ **Equation 13**

Now that we have the trial-by-trial estimated value distributions (Z), we apply CVaR level η to read out values as input to the softmax decision-making function. CVaR can be used to read out a part of either the lower [6] or the upper end [7] of a distribution. Reading out the lowest generates the lowest value while reading out the highest end gives the highest value. Here we set the CVaR levels η from -0.95 to 0.95, with -0.95 reading out 5% lower end of a distribution and 0.95 reading out the top 5% of a distribution, while 0 reading out the mean of the whole distribution. To do this, we first calculated the cumulative distribution function (CDF) of a distribution (Z) (Equation 14) and then find the corresponding percentile of the distribution, denoted as Value at Risk (VaR), for an $\alpha$ level (Equation 15). The CVaR is derived as either the mean of the distribution below the VaR (if η ≤ 0) or the mean of the distribution higher than the VaR (if η > 0) (Equation 16). CVaRs for the two options were put into a softmax function (**Error! Reference source not found.**) to estimate the probability of choosing o ption a.

$$F(Z) = P(Z \leqslant z) \quad \textbf{Equation 14}$$

$$VaR_{\eta}(Z) = \begin{cases} \min\{Z \mid F(Z) \geqslant 1 + \eta\}, \text{ if } -1 < \eta \leqslant 0 \\ \min\{Z \mid F(Z) \geqslant \eta\}, \text{ if } 0 < \eta < 1 \end{cases} \quad \textbf{Equation 15}$$

$$CVaR_{\eta}(Z) = \begin{cases} E\left[Z \mid Z \leqslant VaR_{\eta}(Z)\right], \text{ if } -1 < \eta \leqslant 0 \\ E\left[Z \mid Z \geqslant VaR_{\eta}(Z)\right], \text{ if } 0 < \eta < 1 \end{cases} \quad \textbf{Equation 16}$$

#### Model fitting

All models were fitted in MATLAB using a variational Bayes approach. Only behaviour data from the four same-mean blocks were used for the model fitting. All trials from the same-mean blocks were used in the modelling fitting. Akaike information criterion (AIC) was calculated for each model using the best-fitted parameters for each participant (Equation 17 & Equation 18). $\hat{L}$ denotes the maximized value of the likelihood function of the model M, x: the observed data, k: the number of free parameters in the model. AIC scores were summed across participants, with lower sum AIC indicating better model fit. Delta AIC for each model was calculated by subtracting the AIC score of the best-fitting model in each experiment.

$$AIC = 2 * k - 2\ln(\hat{L}) \quad \textbf{Equation 17}$$

$$\hat{L} = p(x \mid \hat{\theta}, M) \quad \textbf{Equation 18}$$

##### Structural equation model

Structural equation model (SEM) enables assessment of hypothetical causal relations among several (more than two) continuous variables based on their covariance with one another [8, 9]. All structural equation modelling was conducted in Latent Variable Analysis (lavaan) package v.0.6-17 using Maximum Likelihood estimation [10].

Based on the interaction of the ROIs (Hb, VTA, NAcc and mPFC) shown in animal studies [11], we constructed the hypothesized relationships amongst the 4 ROIs as shown in **Fig.S 6**a, i.e. Hb to VTA and then VTA to NAcc and mPFC. We then permute ROI positions within this structure, which results in 16 combinations. Matrixes comprising the filtered time-series of BOLD signals in each ROI from all blocks were used. Akaike information criterion (AIC) was used for model comparison. Delta AIC for each model was calculated by subtracting the AIC score of the winning model for each combination. The winning model was the one with the lowest AICs of all 16 combinations. In **Fig.S 6**b, delta AICs were plotted with x labels corresponding to the starting ROI in this structure.

#### fMRI data analyses

**GLM3:** BOLD=β0+ β1*Response at choice phase + β2*reaction time at choice phase+ β3*mean of choice phase + β4* mean of the first feedback + β5* win amount for the win trials at 2nd feedback + β6* mean for the win trials at 2nd feedback + β7* loss amount for the win trials at 2nd feedback + β8* mean for the loss trials at 2nd feedback + β9* mean of the draw trials + β10* mean of the 3nd feedback (overall outcome feedback).

##### Representational Similarity Analysis (RSA)

RSA was performed using an RSA toolbox adapted for FSL data [2]. For every voxel in each of the ROI (Hb, VTA, NAcc, mPFC) and for each participant, we extracted parameter estimates from the second-level images of the **Error! Reference source not found.** analysis for the mean of the trials for the narrower/narrower option when it was chosen and when its outcome revealed, i.e. $\beta_7$ and $\beta_9$. Then for each ROI, correlations between the voxel-based betas were calculated for each participant. We then ran one-sample t-tests to see whether the correlation coefficients were significantly different from zero across all participants.
